# Inferring synteny between genome assemblies: a systematic evaluation

**DOI:** 10.1101/149989

**Authors:** Dang Liu, Martin Hunt, Isheng. J. Tsai

## Abstract

Identification of synteny between genomes of closely related species is an important aspect of comparative genomics. However, it is unknown to what extent draft assemblies lead to errors in such analysis. To investigate this, we fragmented genome assemblies of model nematodes to various extents and conducted synteny identification and downstream analysis. We first show that synteny between species can be underestimated up to 40% and find disagreements between popular tools that infer synteny blocks. This inconsistency and further demonstration of erroneous gene ontology enrichment tests throws into question the robustness of previous synteny analysis when gold standard genome sequences remain limited. In addition, determining the true evolutionary relationship is compromised by assembly improvement using a reference guided approach with a closely related species. Annotation quality, however, has minimal effect on synteny if the assembled genome is highly contiguous. Our results highlight the need for gold standard genome assemblies for synteny identification and accurate downstream analysis.

**Author summary:** Genome assemblies across all domains of life are currently produced routinely. Initial analysis of any new genome usually includes annotation and comparative genomics. Synteny provides a framework in which conservation of homologous genes and gene order is identified between genomes of different species. The availability of human and mouse genomes paved the way for algorithm development in large-scale synteny mapping, which eventually became an integral part of comparative genomics. Synteny analysis is regularly performed on assembled sequences that are fragmented, neglecting the fact that most methods were developed using complete genomes. Here, we systematically evaluate this interplay by inferring synteny in genome assemblies with different degrees of contiguation. As expected, our investigation reveals that assembly quality can drastically affect synteny analysis, from the initial synteny identification to downstream analysis. Importantly, we found that improving a fragmented assembly using synteny with the genome of a related species can be dangerous, as this *a priori* assumes a potentially false evolutionary relationship between the species. The results presented here re-emphasize the importance of gold standard genomes to the science community, and should be achieved given the current progress in sequencing technology.

## Introduction

The essence of comparative genomics lies in how we compare genomes to reveal species’ evolutionary relationships. Advances in sequencing technologies have enabled the generation and exploration of many new genomes across all domains of life [1–8]. Unfortunately, in most instances correctly aligning even just two genomes at base-pair resolution can be challenging. A genome usually contains millions or billions of nucleotides and is different from the genome of a closely related species as a result of evolutionary processes such as mutations, chromosomal rearrangements, and gene family expansion or loss. There are computational complexities when trying to align and assign multiple copies of DNA, such as transposable elements that are identical to each other [9–12]. In addition, it has been shown that popular alignment methods disagree with each other [9].

An alternative and arguably more practical approach relies on the identification of synteny blocks [13,14], first described as homologous genetic loci presenting on the same chromosome [15,16]. Currently it is more formally defined as regions of chromosomes between genomes that share a common order of homologous genes derived from a common ancestor [17,18]. Alternative names such as conserved synteny or collinearity have been used interchangeably [13,19–22]. Comparisons of genome synteny between and within species have provided an opportunity to study evolutionary processes that lead to diversity of chromosome number and structure in many lineages across the tree of life [23,24]; early discoveries using such approaches include chromosomal conserved regions in nematodes and yeast [25–27], evolutionary history and phenotypic traits of extremely conserved Hox gene clusters across animals and MADS-box gene family in plants [28,29], and karyotype evolution in mammals [30] and plants [31]. Analysis of gene synteny against closely related species is now the norm in every new published genome. However, assembly quality comes into question as it has been demonstrated to affect subsequent analysis such as annotation or rate of lateral transfer [32,33].

In general, synteny identification is a filtering and organizing process of all local similarities between genome sequences into a coherent global picture [34]. Orthologous relationships of protein-coding genes are used as anchors to position statistically significant local alignments. Approaches include the use of a directed acyclic graph [35,36], a gene homology matrix (GHM) [37], and an algorithm using reciprocal best hits (RBH) [38]. All of these methods generally agree on long synteny blocks, but have differences in handling local shuffles as well as the resolution of synteny identification [34,38]. Better resolution of micro-rearrangements in synteny patterns has been shown when using an improved draft genome of *Caenorhabditis briggsae* versus *Caenorhabditis elegans* [26,39]. Hence, synteny analysis depends highly on assembly quality. For example, missing sequences in assembly lead to missing gene annotations and subsequently missing orthologous relationship [40]. With respect to assembly contiguation, it still remains unclear whether assembly fragmentation affects the homology assignments for deciding anchors, features of genes for examining order and gaps, or other factors in synteny analysis.

In this study, we focus on how assembly quality affects the identification of genome synteny. We investigate the correlation between error rate (%) in detecting synteny and the level of assembly contiguation using four popular software packages (DAGchainer [35], i-ADHoRe [37], MCScanX [36], and SynChro [38]) on four nematodes: *Caenorhabditis elegans, Caenorhabditis briggsae*, *Strongyloides ratti* and *Strongyloides stercoralis*. We also carried out and explored the effects of three scenarios associated with synteny analysis: gene ontology (GO) enrichment, reference-guided assembly improvement, and annotation quality. Our findings show that assembly quality does matter in synteny analysis, and fragmented assemblies ultimately lead to erroneous findings. In addition, the true evolutionary relationship may be lost if a fragmented assembly is improved using a reference-guided approach. Our main aim here is to determine a minimum contiguation of assembly for subsequent synteny analysis to be trustworthy, which should be possible using the latest sequencing technologies [41].

## Results

### Definition of synteny block, break and coverage

We begin with some terminology that is used throughout this study. As shown in Fig 1, a synteny block is defined as a region of genome sequence spanning a number of genes that are orthologous and co-arranged with another genome. Orientation is not considered (Fig 1, Block a and b). The minimum number of co-arranged orthologs which are said to be the anchors can be set and vary between different studies. A larger number of minimum anchors may result in fewer false positives, but also a more conservative estimate of synteny blocks (S1 Fig). In some programs, some degrees of gaps are tolerated (Fig 1, Block c), which are defined as the number of skipped genes or the length of unaligned nucleotides. A score is usually calculated, and synteny breaks are therefore regions that do not satisfy a certain score threshold. Possible scenarios that lead to synteny breaks include a lack of anchors in the first place (Fig 1, break a), a break in anchor order (Fig 1, break b), or gaps (Fig 1, break c). Genome insertions and duplications are potential causes of over-sized gaps. An example is Break c of Fig 1, which is due to either a large unaligned region (Fig 1, P^1^-Q^1^ and Q^2^-R^2^) or a high number of skipped genes (Fig 1, S^2^-T^2^-X^2^ within Q^2^-R^2^). Alternatively, an inversion (Fig 1, orthologs K and L), deletion or translocation (Fig 1, ortholog X) may cause a loss of anchors (Fig 1, gene D in species 1) or a break in the arrangement of anchors. Typically, synteny coverage is commonly used as a summary metric, which is obtained by dividing summed length of blocks by genome size. Note that synteny coverage is asymmetrical, as demonstrated by the difference of block length in Block c (Fig 1).

**Fig 1.**
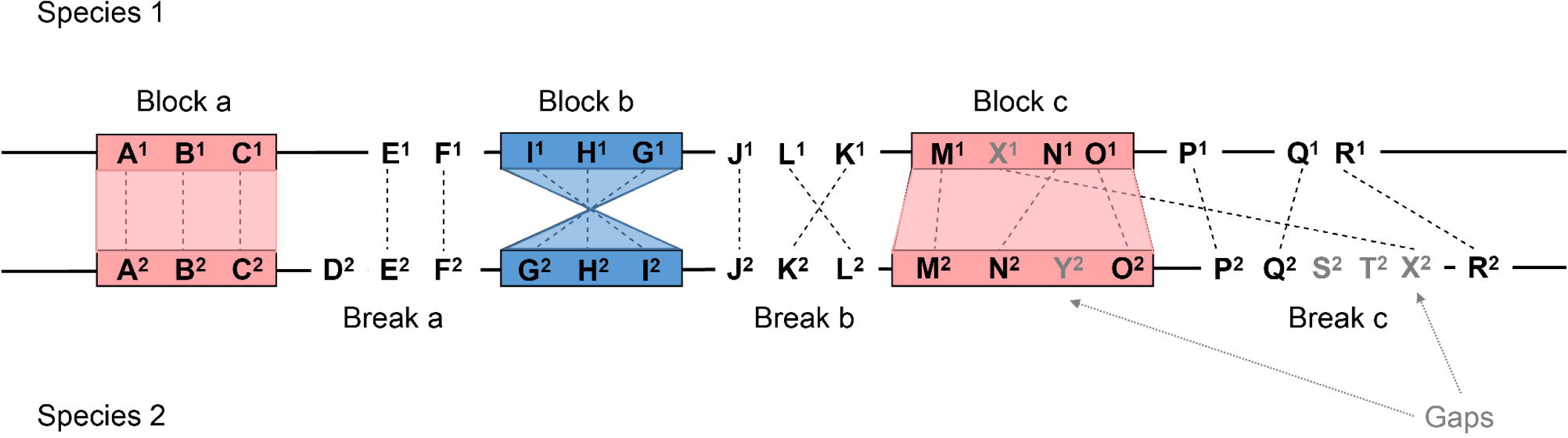
Definition of synteny block and synteny break. Genes located on chromosomes of two species are denoted in letters. Each gene is associated with a number representing the species they belong to (species 1 or 2). Orthologous genes are connected by dashed lines and genes without an orthologous relationship are treated as gaps in synteny programs. Under the criteria of at least three orthologous genes (anchors): a synteny block can be orthologs with the same order (Block a), reverse order (Block b), or allowing some gaps (Block c). In contrast, cases of causing a synteny break can be lack of orthologs (Break a), gene order (Break b) or gaps (Break c).

### Evaluation of synteny identification programs in fragmented assemblies

There are several programs developed to identify synteny blocks, which can produce quite different results [14]. Our first aim is to systematically assess the synteny identification of four popular tools: DAGchainer [35], i-ADHoRe [37], MCScanX [36], and SynChro [38]. As whole genome alignments between bacteria, which have relatively small genomes, is becoming common practice [42], we conduct this study on species with larger genome sizes. We chose *Caenorhabditis elegans*, a model eukaryote with a 100 megabase (Mb) reference genome. Our first question was whether these programs would produce 100% synteny coverage if the *C. elegans* genome was compared to itself. As expected, all tools accurately achieved almost 100% synteny coverage (Fig 2 and S2 Fig).

**Fig 2.**
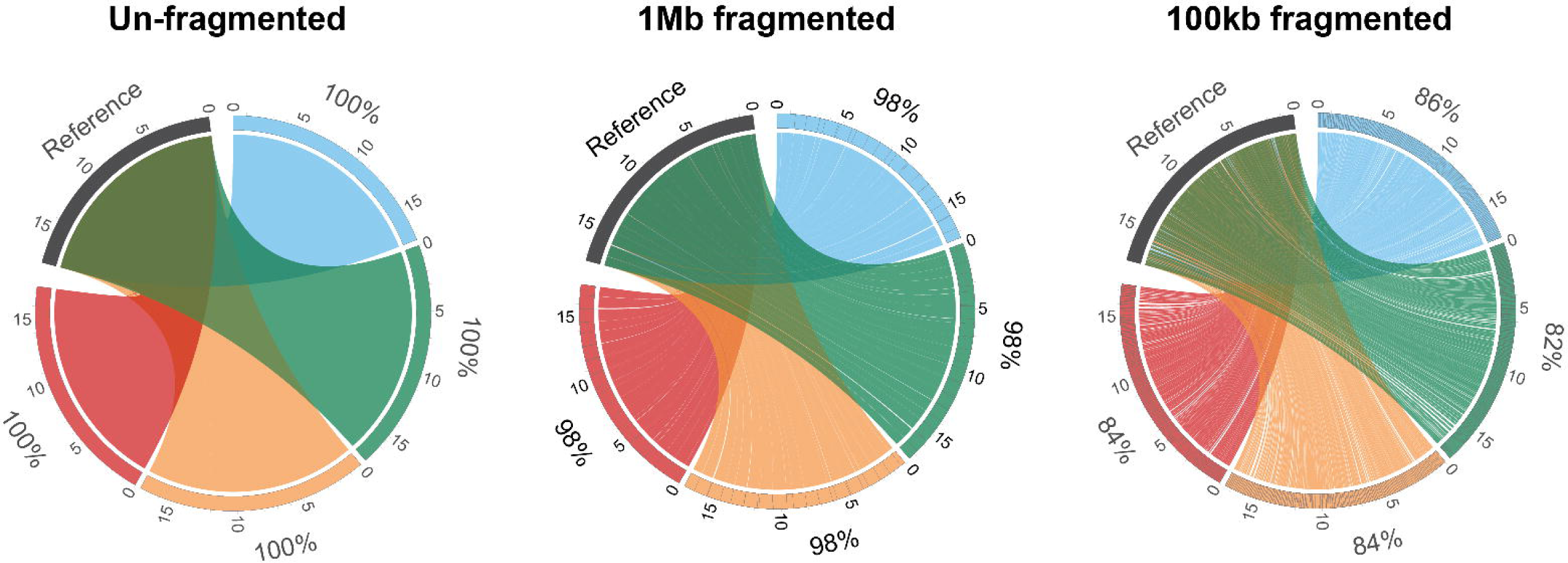
Synteny blocks identification between *C. elegans* chromosome IV. The original sequence is used as the reference and coloured in black. Established synteny regions (outer number stands for synteny coverage) of the 4 different program packages: DAGchainer (red), i-ADHoRe (yellow), MCScanX (green) and SynChro (light blue) are joined to query sequences with different levels of fragmentation (un-fragmented, 1Mb and 100kb fragmented). Chromosome positions are labeled in megabases (Mb). For plots of other chromosomes see S1-3 Figs.

We then fragmented the *C. elegans* genome into fixed intervals of either 100kb, 200kb, 500kb or 1Mb to evaluate the performance of different programs when using self-comparisons (Methods). Synteny coverages of the fragmented assembly (query) against the original assembly (reference) were calculated for both query and reference sequences. As expected, synteny coverage decreased as the assembly was broken into smaller pieces. For example, an average of 16% decrease in synteny coverage was obtained using the assembly with fixed fragment size of 100kb (S2 Table). Sites of fragmentation are highly correlated with synteny breaks (Fig 2). One explanation is that fragmented assembly introduced breaks within genes resulting in loss of anchors (Fig 1, Break a), which can be common in real assemblies if introns contain hard to assemble sequences [32]. Another explanation is that the breaks between genes lead to the number anchors not reaching the required minimum (Fig 1, Break a). To assess our fragmented approach to real data, we obtained a recent publicly available genome of *C. elegans* using long reads data (Methods). The assembly has an N50 of ~1.6Mb and we annotated this assembly *de novo*. A synteny coverage of 98.9% was obtained which is very similar to our 1Mb fragmented assemblies of 98.4% (S2 Fig and S1 Table) suggesting robustness in our fragmentation approach.

More fragmented assemblies led to greater differences in synteny coverage predicted between the four tools (Fig 2 and S2-4 Figs). We carefully examined regions where synteny was predicted in some programs but not the other (Figs 2 and 3). Fig 3 demonstrates such a case of disagreement. DAGchainer and i-ADHoRe produced the same results, whilst MCSanX and SynChro detected less and more synteny, respectively (Fig 3). MCSanX’s gap scoring scheme used a stricter synteny definition, and more synteny blocks can be identified when the gap threshold was lowered (Fig 3, situation a; also see S5 Fig). SynChro has its own dedicated orthology assignment approach that assigns more homologous anchors (Fig 3, situation b). Additionally, SynChro uses only 2 genes as anchors to form a synteny block, while the default is at least five gene anchors in other three tools (Fig 3, situation b). Together, these results suggest that the default parameters set by different tools will lead to differences in synteny identification and need to be tuned before undertaking subsequent analysis.

**Fig 3.**
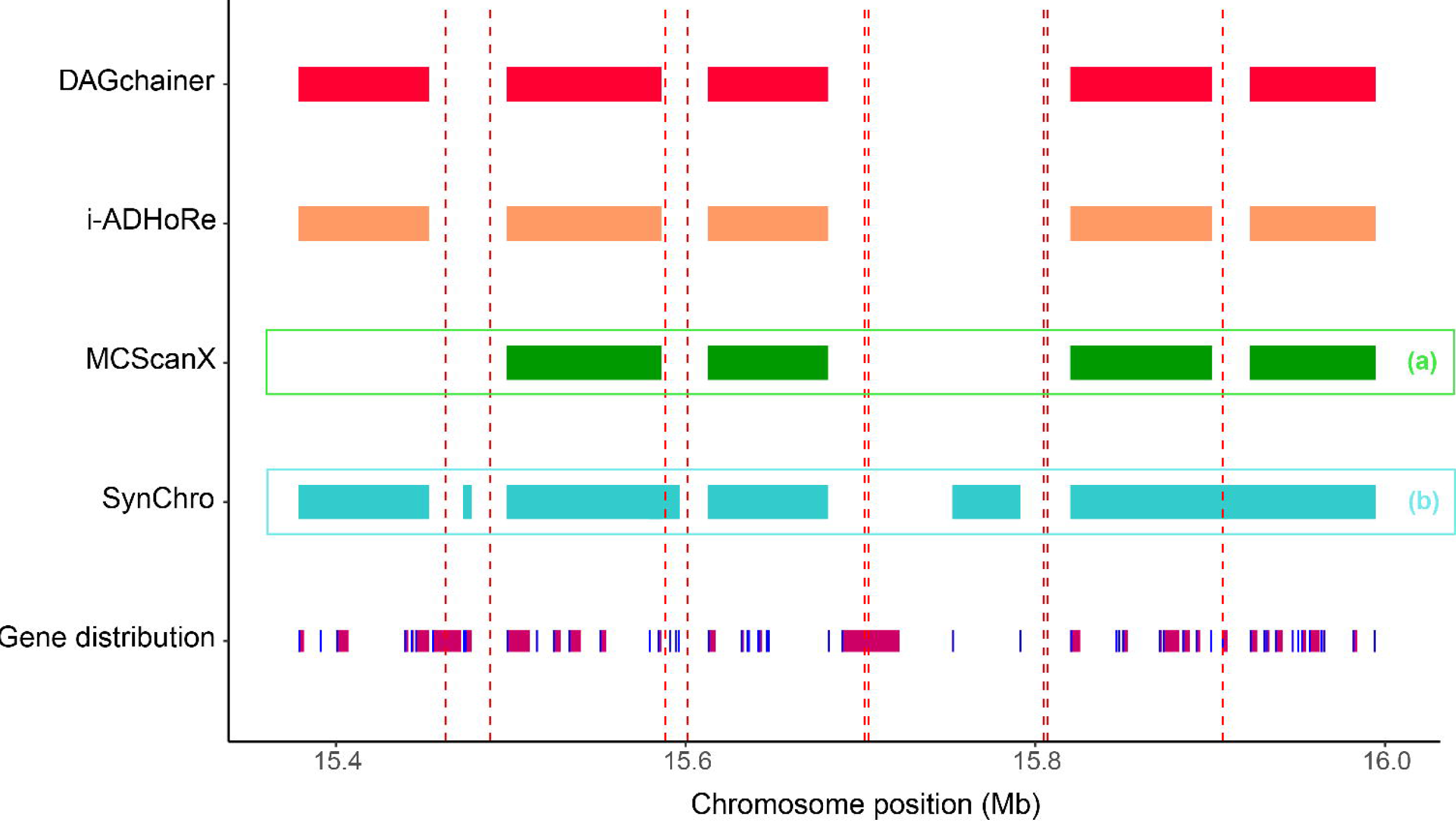
A zoomed-in 600kb region of synteny identification between the reference *C. elegans genome* and a 100kb fragmented assembly. Synteny blocks defined by the four detection programs DAGchainer (red), i-ADHoRe (yellow), MCScanX (green) and SynChro (light blue) are drawn as rectangles. Fragmented sites are labeled by vertical red dashed lines. Genes are shown as burgundy rectangles, with gene starts marked using dark blue lines. Two scenarios are marked: a) synteny block was not identified by MCSanX, and b) synteny blocks only detected by SynChro.

### Contribution of assembly contiguation and intrinsic species effect to synteny analysis

To quantify the effect of assembly contiguation in synteny analysis, we used four nematode genomes: *Caenorhabditis elegans, Caenorhabditis briggsae*, *Strongyloides ratti* and *Strongyloides stercoralis*. Nematodes are useful models in synteny analysis as 1) extensive chromosomal rearrangement are hallmarks of their genome evolution [7,25,26,43,44]; 2) the genome sequences are highly contiguous and assembled into chromosomes [7,25,26,43]. Also, these two genera were chosen to investigate the intrinsic species effect as they differ in their density of genes (Table 1). Our fragmentation approach was first used to break the *C. elegans* and *S. ratti* genomes into fixed sequence size of either 100kb, 200kb, 500kb, or 1Mb. Here, we define the error rate as the difference between the original synteny coverage (almost at 100%) and when assembly is fragmented. For each fixed length, the fragmentation was repeated 100 times so assemblies are broken at different places to obtain a distribution. As expected, there is a positive correlation between error rate and level of fragmentation. Specifically, the median error rate can be as high as 18% at assemblies with sequence length of 100kb (S2 Table). Amongst the four tools, fragmented assemblies have largest effect in MCScanX and least in SynChro (Fig 4 and S2 Table).

**Fig 4.**
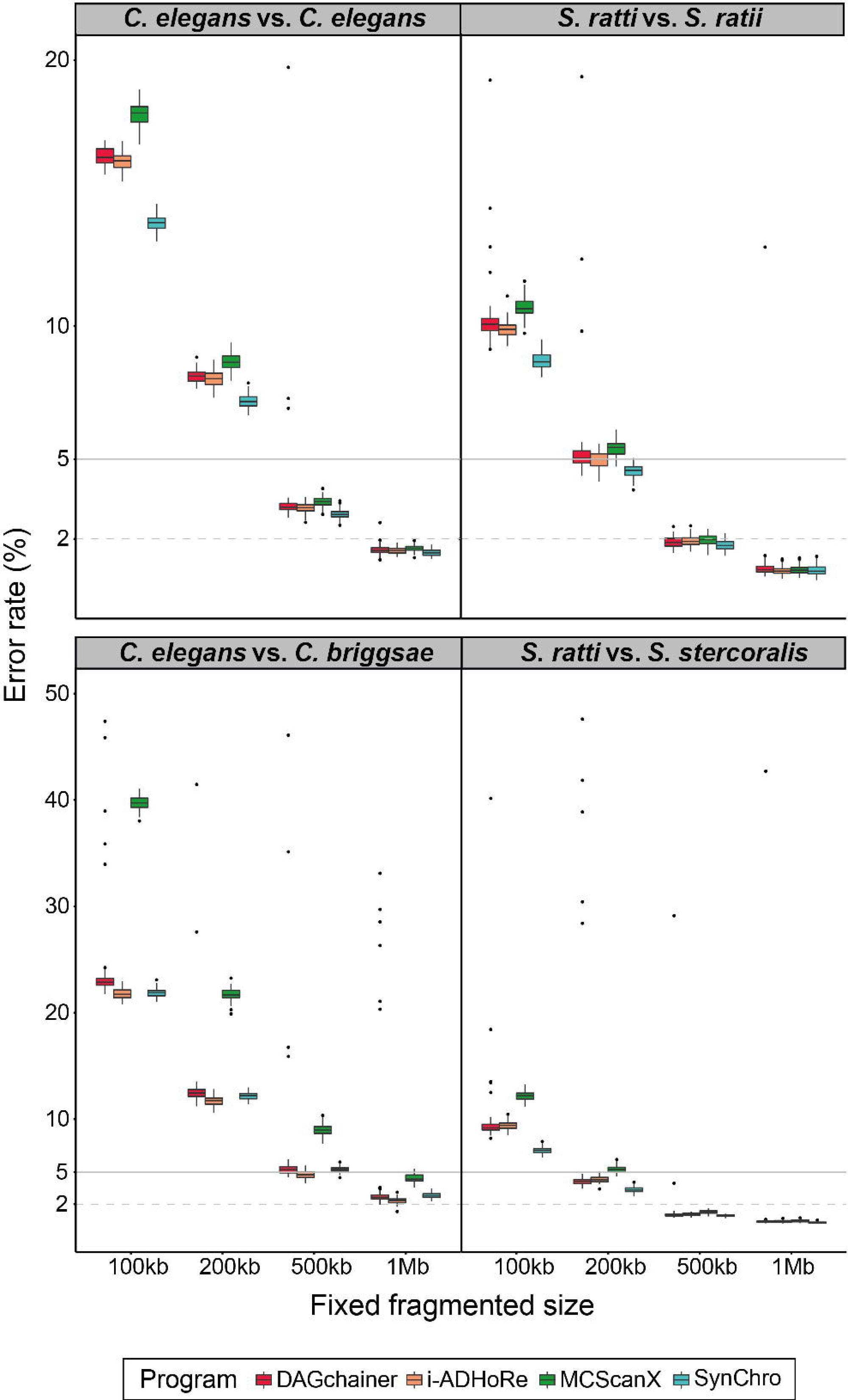
Error rate (%) of synteny identification in fragmented assemblies. The error rate is defined as the difference between the synteny coverage calculated with original genome (almost 100%) and that in fragmented assemblies, where in both cases the original assembly was used as the reference. 5 % and 2 % error rate positions are marked by grey solid and dashed lines, respectively. Different pairs of synteny identification are separated in different panels. The upper panels are self-comparisons, while the bottom are comparisons between closely related species. Note that for a clear visualization of pattern changes, the scales of error rate are different between upper and bottom panels. Colors represent different types of synteny detection programs.

**Table 1.** Genomic features of *Caenorhabditis* and *Strongyloides* species.

|  | *C. elegans* | *C. briggsae* | *S. ratti* | *S. stercoralis* |
| --- | --- | --- | --- | --- |
| Genome size (Mb) | 100.3 | 108.4 | 43.2 | 42.7 |
| Number of genes | 20,247 | 21,814 | 12,453 | 13,098 |
| Gene density (genes/Mb) | 201.9 | 201.3 | 288.6 | 307.0 |
| Gene coverage (%) | 63.1 | 59.7 | 50.9 | 51.8 |
| Median gene length (bp) | 1,972 | 1,964 | 1,281 | 1,195 |
| Mean gene length (bp) | 3,124.2 | 2,967.5 | 1,763.8 | 1,687.8 |
| Median intergenic length | 925 | 1,183 | 923 | 808 |
| Mean intergenic length | 2,209.5 | 2,394.6 | 1,712.0 | 1,513.8 |

Features that may play a key role in synteny detection are highlighted in yellow.

A common analysis when reporting a new genome is inferring synteny against a closely related species. Hence, we reanalyzed synteny between *C. elegans* and *C. briggsae*, *S. ratti* and *S. stercoralis.* On average, the four tools found 77% and 83% synteny between *C. elegans*-*C. briggsae* and *S. ratti*-*S. stercoralis* respectively (S2 Table). In contrast to broad agreement on within-species self-comparisons, the tools varied considerably on these inter-species (i.e. more diverged) comparisons (S6 Fig and S2 Table). For example, in the *C. elegans*-*C. briggsae* comparisons, a difference of 25% in synteny coverage was found between the results of i-ADHoRe and SynChro (S6 Fig and S2 Table), while this tool variation was interestingly much lower in *S. ratti*-*S. stercoralis* with only 9% difference (S2 Table). To increase the complexity, we fragmented *C. briggsae* and *S. stercoralis* into fixed sequence sizes using the same approach as previously mentioned and compared them with the genome of *C. elegans* and *S. ratti*, respectively. We found that MCScanX still underestimated synteny the most as the scaffold size decreased from 1Mb to 100 kb. Strikingly, the median error rate was high as 40% in *C. elegans-C. briggsae* but only 12% in *S. ratti*-*S. stercoralis* comparisons (Fig 4). This observation suggests that higher gene density leads to more robustness in synteny detection in fragmented assemblies as more anchors (genes) are available in a given sequence (Table 1 and S2 Table).

### Erroneous findings using fragmented assemblies in synteny analysis

Functional enrichment of genes of interest are usually investigated after the establishment of orthology and synteny [26,45–48]. Synteny breaks contain rearrangements, novel genes, or genes that are too diverged to establish an orthologous relationship or have undergone expansion or loss. Functions of these genes are often of interest in comparative genomics analyses. To further estimate the effect of poor assembly contiguation on synteny analysis, GO enrichment was performed for genes present in synteny breaks in the original assembly of *C. elegans* versus fragmented assemblies of *C. briggsae*. This approach was then repeated 100 times each with assemblies fragmented randomly. We found that fragmented assemblies lead to GO terms originally not in the top 100 ranks then consistently appearing in the top 10 during the 100 replicates (Fig 5 and S3-6 Table). Furthermore, the orders of the original top 10 GO terms shifted in fragmented assemblies (Fig 5 and S3-6 Table). In addition, the 10th top GO term failed to appear in the top 10 in 100 replicates of 100kb and 200kb fragmented assemblies (Fig 5 and S3-6 table). These results suggest that an underestimation of synteny relationship due to poor assembly contiguation can lead to a number of erroneous findings in subsequent analysis.

**Fig 5.**
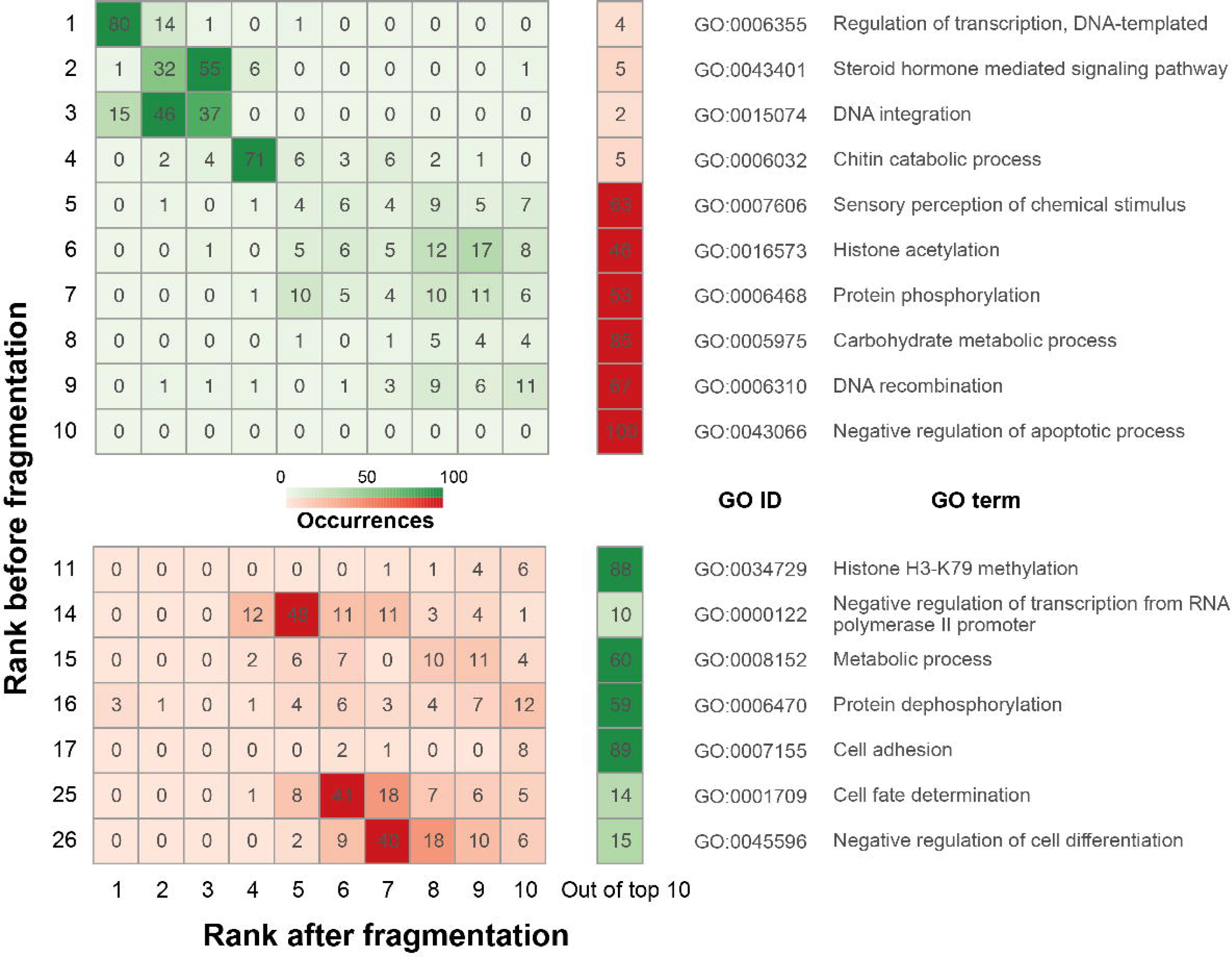
Comparison of gene ontology (GO) enriched terms in *C. briggsae* synteny break between *C. elegans* vs. *C. briggsae* and 100 replicates of *C. elegans* vs. 100kb fragmented *C. briggsae*. Top ranks of GO terms in the original comparison are shown in the Y axis. For original top ranking GO terms, only those that appeared more than 10 times in top 10 ranks of after-fragmentation comparison replicates were displayed (see S6 Table for more details). The X axis shows top 10 ranks and rank “out of top 10” in the comparison when assemblies were fragmented. The darkness of color is proportional to the occurrence of the GO term in that rank within 100 replicates. Regions in red are indications of occurred errors.

### True synteny is compromised by reference-guided assembly methods

Although assembly quality plays an important role in synteny analysis, it has been demonstrated that poor assembly contiguity of one species can be scaffolded by establishing synteny with a more contiguous assembly of a closely related species [40] [49–51]. However, we hypothesised that the true synteny relationship between two species may be incorrectly inferred when an assembly of one species is scaffolded based on another closely related species, by assuming the two genomes are syntenic. In order to investigate this, ALLMAPS [51] was used to order and orient sequences of 100kb fragmented *C. briggsae* based on *C. elegans* as well as 100kb fragmented *S. stercoralis* assembly based on *S. ratti*. ALLMAPS improved both fragmented assemblies impressively, increasing the N50 from 100kb to 19Mb and 15Mb in *C. briggsae* and *S. stercoralis*, respectively (S7 Table). Synteny coverage from these improved assemblies was even higher than the original fragmented 100kb sequences in MCScanX, much lower in i-ADHoRe, and similar in DAGchainer and SynChro (Fig 6). In addition, despite synteny coverage close to that of the original assemblies in DAGchainer and SynChro, further investigation of synteny block linkages in *C. elegans*-*C. briggsae* using identification from DAGchainer revealed frequent false ordering and joining of synteny blocks. Intra-chromosomal rearrangements are common between *C. elegans* and *C. briggsae*, but ALLPMAPS improved assembly have shown a false largely collinear relationship in the chromosomes between the two species (Fig 7). Hence, reference guided assembly improvement produces pseudo-high quality assemblies that may have ordering biased towards the reference genome and may not reflect the true evolutionary scenario.

**Fig 6.**
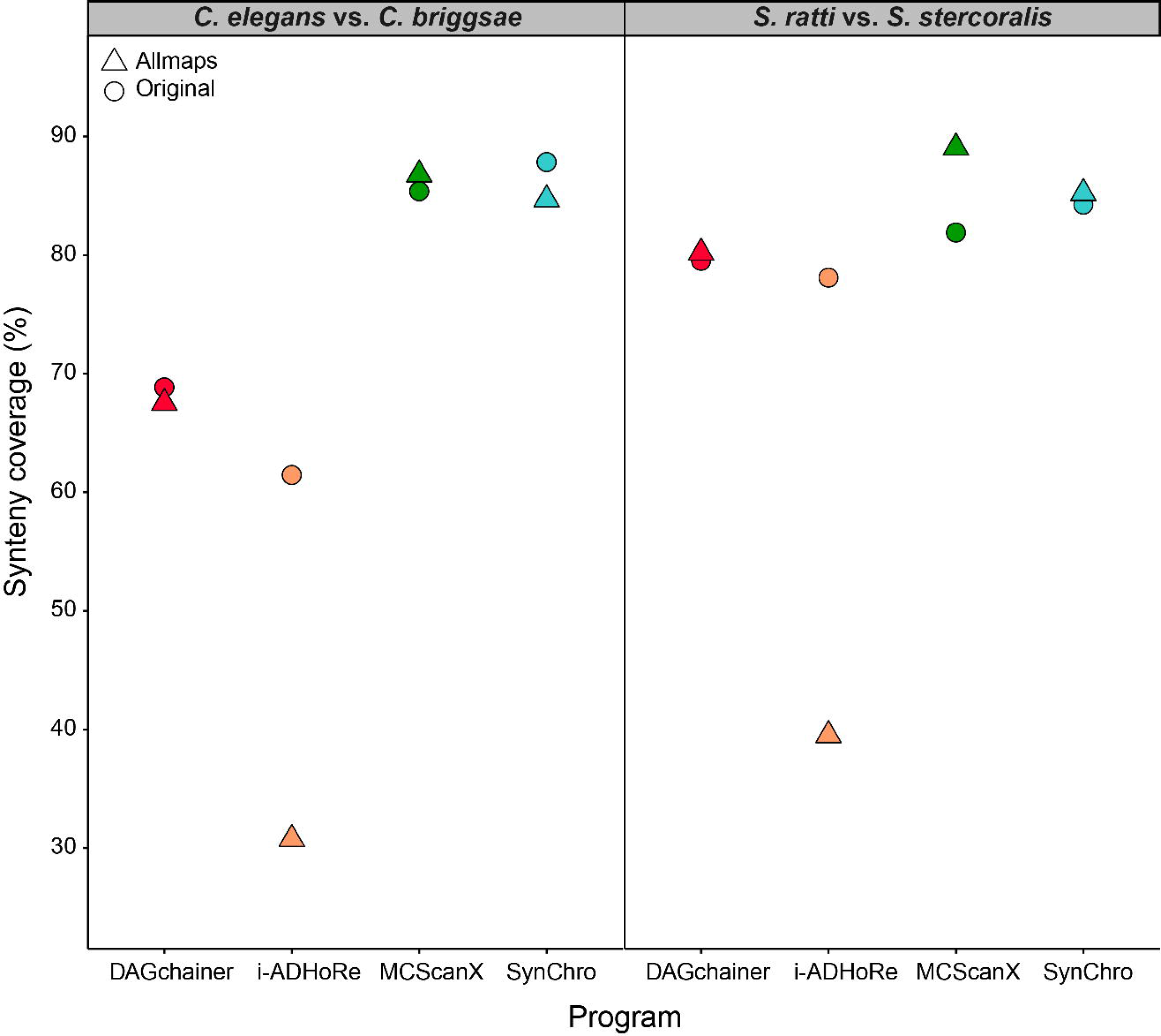
Synteny coverage (%) between *C. elegans* and *S. ratti* assemblies against original or ALLMAPS improved 100kb fragmented *C. briggsae* and *S. stercoralis.*

**Fig 7.**
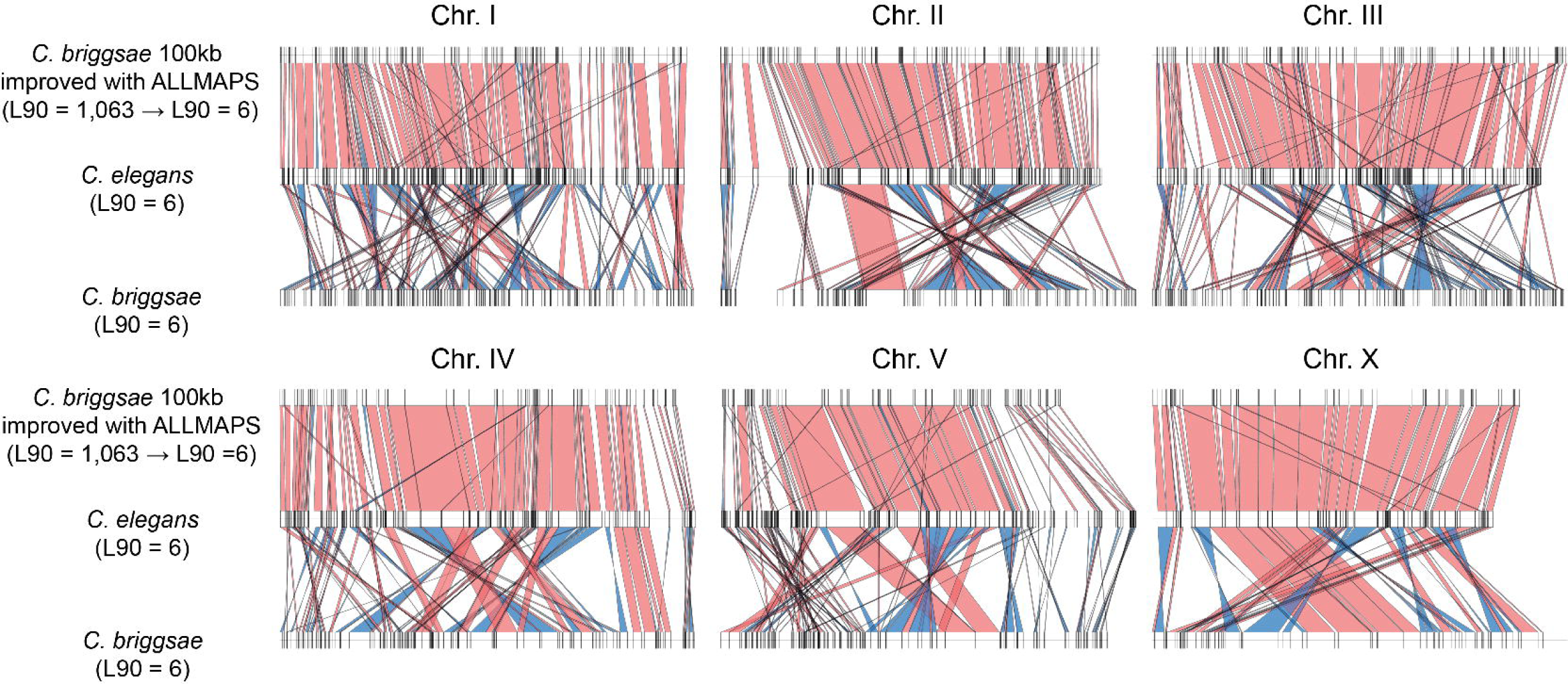
Synteny linkage between *C. elegans* vs. original *C. briggsae* assemblies and *C. elegans* vs. ALLMAPS *C. briggsae* assembly. ALLMAPS assembly with L90 = 1,063 from 100kb fragmented *C. briggsae* assembly with L90 = 6 (top), original *C. elegans* assembly with L90 = 6 (middle) and original *C. briggsae* assembly with L90 = 6 (bottom) are shown in different horizontal lines. Vertical lines on chromosome lines show the start/end positions of the first/last gene in a synteny block. Each panel shows a separate chromosome. Block linkages in the same orientation are labeled in red, while those in inverted orientation are labeled in blue.

### Annotation quality has little effect on synteny analysis

Genome annotation is a bridging step between genome assembly and synteny analysis. An incomplete annotation may lead to lack of homology information in synteny analysis. We compared synteny coverage in three datasets of *C. elegans* that differ in quality of annotation: 1) manually curated WormBase [52] *C. elegans* annotation, 2) optimized Augustus [53] annotation with its built-in *Caenorhabditis* species training set, and 3) semi-automated Augustus annotation with the BUSCO [54] nematoda species training set. In either case, we found that synteny coverage varies at most 0.02% in reference genome (Table 2). As a result, with a well-assembled genome and proper species training set, the quality of annotation has little effect on synteny analysis, compared to assembly quality.

**Table 2.**
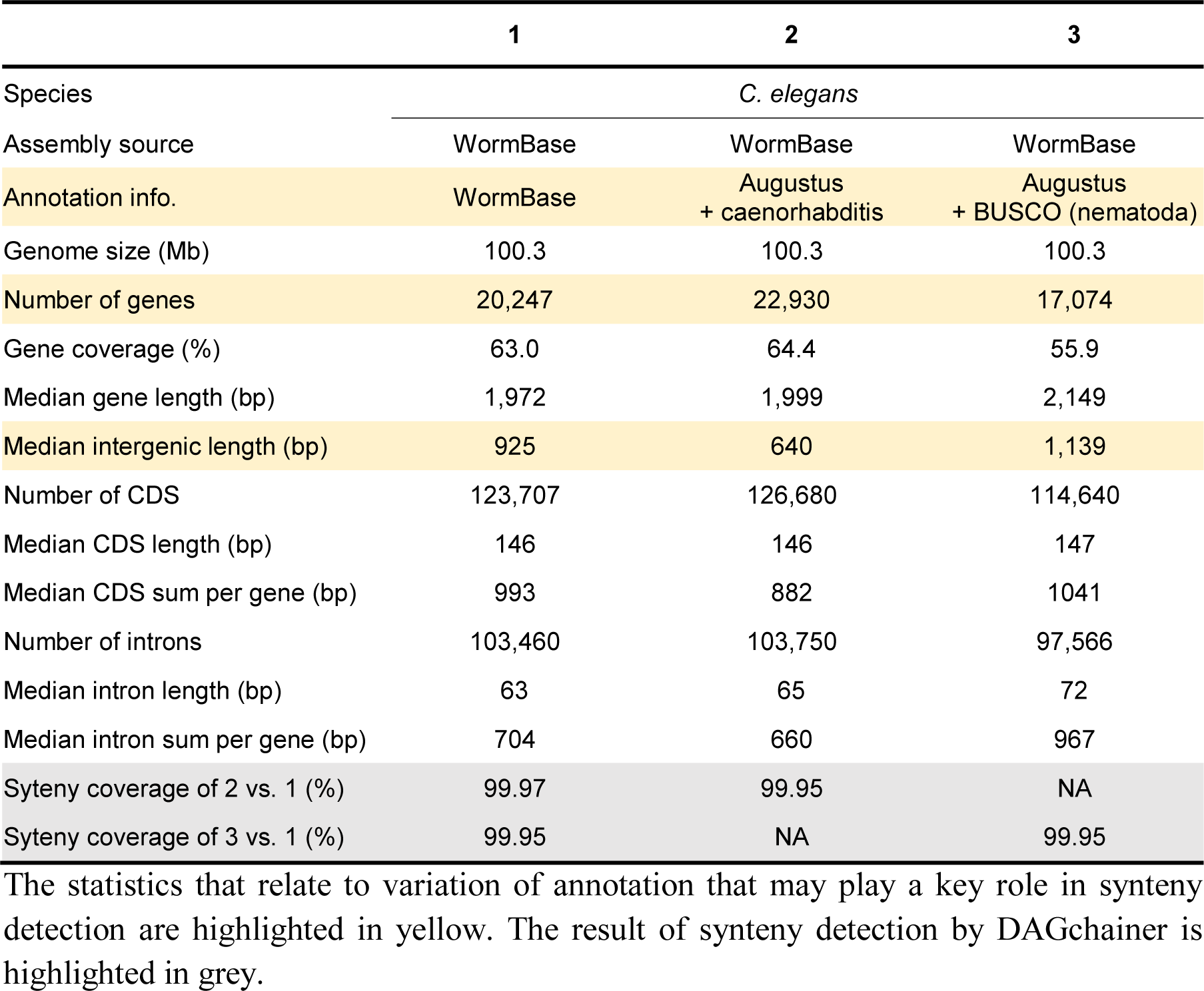
Statistics of *C. elegans* annotations used for synteny analysis

## Discussion

Synteny analysis is a practical way to investigate the evolution of genome structure [28–30,55]. In this study, we have revealed how genome assembly contiguity affects synteny analysis. We present a simple scenario of breaking assembly into more fragmented state, which only mimics part of the poor assembly problem. Our genome fragmentation method randomly breaks sequences into pieces with the same size, which gives rise to an almost even distribution of sequence length. This is a simplification of real assemblies, which usually comprise few large sequences and many more tiny sequences that are difficult to assemble because of their repetitive nature [25,26]. It is probable that we overestimate error rate in regions that can be easily assembled but underestimate error rate in regions that will be more fragmented. Note some of the sequences in real assemblies may contain gaps (scaffolds) which will result in more missing genes and will result in further underestimation of synteny. Our result is quite similar when a *de* novo Pacbio *C. elegans* assembly was compared to the reference genome (S1 Table). The use of long read technology is becoming the norm in *de novo* assembly projects. Assemblies with lower contiguation were not compared here as we emphasize the responsibility of research groups to produce assemblies that are of the higher contiguity made possible by long reads [56].

Synteny identification from different programs (*i.e.*, DAGchainer [35], i-ADHoRe [37], MCScanX [36], and SynChro [38]) performed across different species (*C. elegans*, *C. briggsae*, *S. ratti* and *S. stercoralis*) have allowed us to examine the wide-ranging effects of assembly contiguation on downstream synteny analysis. Although the four programs tend to produce the same results when the original assembly is compared to itself, this was no longer the case as assemblies become fragmented. It is interesting to note that DAGchainer and MCSanX use the same scoring algorithm for determining synteny regions, except that DAGchainer uses the number of genes without orthology assignment as gaps while MCSanX uses unaligned nucleotides to do this. When comparing closely related species, results of the four programs fluctuate even without fragmentation in *Caenorhabditis* species, while the pattern remains similar to self-comparison in *Strongyloides* species. Sensitivity of synteny identification drops sharply as the genome assembly becomes fragmented, and thus genome assembly contiguation must be considered when inferring synteny relationships between species. Our fragmentation approach only affects N50, which mostly leads to loss of anchors in synteny analysis. Other scenarios such as unknown sequences (NNNs) in the assembly, or rearrangements causing a break in anchor ordering (Fig 1, break b), or insertions/deletions (Fig 1, break c) were not addressed and may lead to greater inaccuracies.

We have shown that genomic features such as gene density and length of intergenic regions play an essential role during the process of synteny identification (Fig 4 and Table 1; S2 Table). Synteny identification can be established more readily in species with higher gene density or shorter intergenic space, which is related to the initial setting of minimum anchors for synteny identification (Fig 1 and S1 Fig). Repetitiveness of paralogs is another factor in how anchors were chosen from homology assignment. For example, we found that synteny coverage is low along chromosomal arm regions of *C. elegans* in both self-comparison and versus *C. briggsae*, which has been reported to have expansion of G protein-coupled receptor gene families [25] (Fig 2 and S6 Fig). Nevertheless, this case may be a result of a combination of repetitive paralogs and high gene density.

Interestingly, synteny comparison with improved assemblies using ALLMAPS [51] exhibited unexpected variation among programs. Unfortunately, we did not resolve the reason behind sharp decrease of synteny coverage in i-ADHoRe (Fig 6). Nevertheless, we have shown that it is dangerous to improve an assembly using a reference from closely related species without *a priori* information about their synteny relationship. Subsequent synteny identification would be misleading if the same reference was compared again. We also considered the interplay between genome annotation, assembly and synteny identification. Although it may be intuitive to assume lower annotation quality can lead to errors in synteny analysis, we demonstrated that such influence was minimal if an initial genome assembly of good contiguation is available (Table 2).

In conclusion, this study has demonstrated that a minimum quality of genome assembly is essential to synteny analysis. As a recommendation, to keep the error rate below 5% in synteny identification, we suggest an N50 of 200kb and 1Mb is required when gene density of species of interest are 290 and 200 genes per Mb, respectively (Table 1 and S2 Table). This is a minimum standard and a higher N50 may be required for other species with lower gene density or highly expanded gene families.

## Materials and Methods

### Data Preparation and Fragmentation Simulation

Assemblies and annotations of *C. elegans* and *C. briggsae* (release WS255), *S. ratti* and *S. stercoralis* (release WBPS8) were obtained from WormBase (http://www.wormbase.org/) [24]. A new assembly of *C. elegans* using long reads was obtained from a Pacific Bioscienceces Dataset (https://github.com/PacificBiosciences/DevNet). Since some genes produce multiple alternative splicing isoforms and all of these isoforms represent one gene (locus), we used the longest isoform as a representative. Further, non-coding genes were also filtered out from our analysis. To simulate the fragmented state of assemblies, a script was made to randomly break scaffolds into fixed size of fragments (https://github.com/dangliu/Assembly-breaking.git). Sequences shorter than the fixed length were kept unchanged.

**Fig 8.**
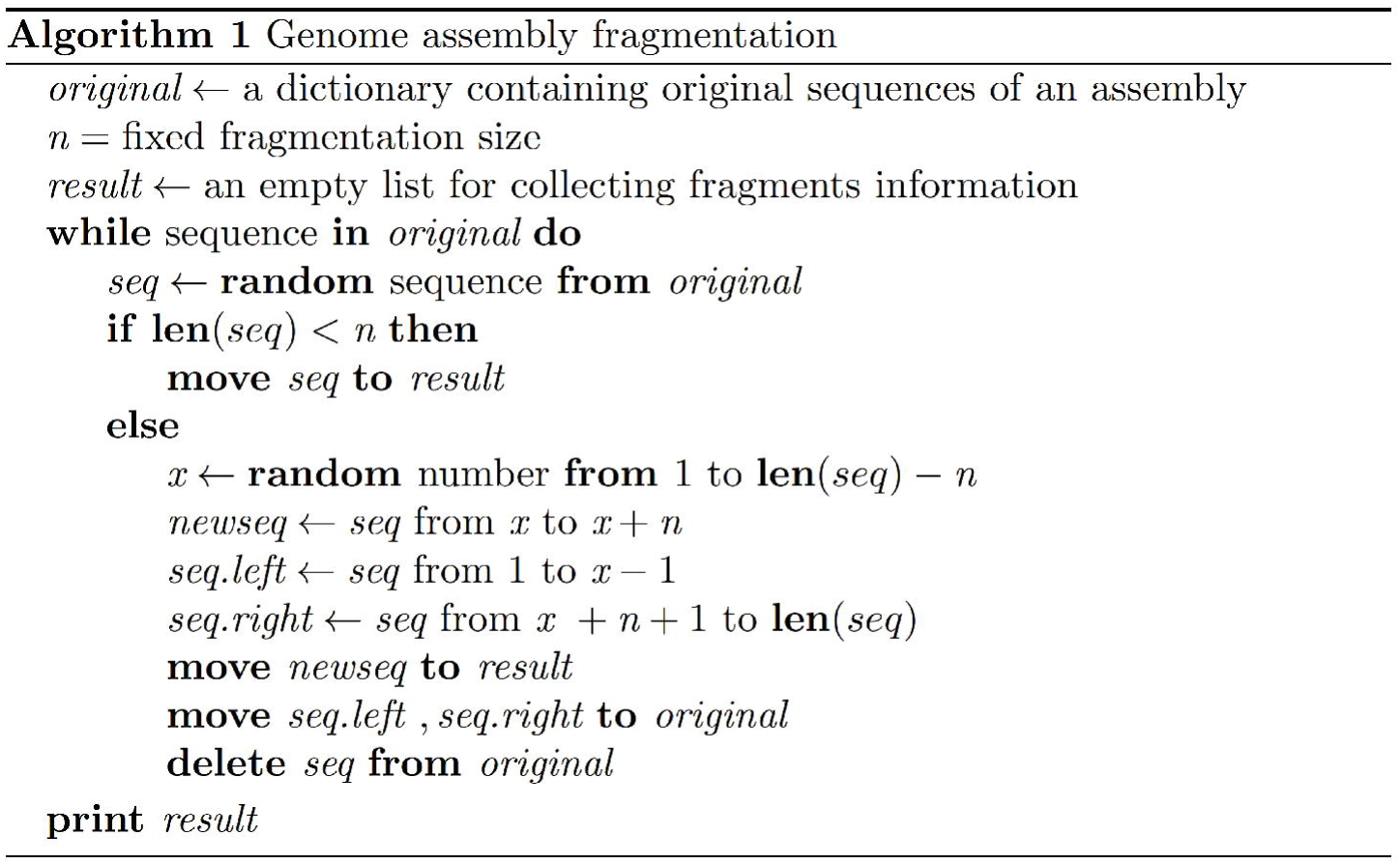
Pseudocode of genome assembly fragmentation.

### Identification of Synteny Blocks

Four different programs were used to identify synteny blocks: DAGchainer [35], i-ADHoRe [37] (v3.0), MCScanX [36] and SynChro [38]. Settings for each program were modified to resemble each other. All of the programs use gene orthology to find anchor points during process of synteny blocks detection. For DAGchainer, i-ADHoRe and MCScanX, we obtained gene orthology from OrthoFinder [57] (v0.2.8). For SynChro, it has an implemented program called OPSCAN to achieve scanning of gene orthology. We arranged the parameters, DAGchainer (accessory script filter_repetitive_matches.pl was run with option 5 before synteny identification as recommended by manual; options: -Z 12 -D 10 -A 5 -g 1), i-ADHoRe (only top 1 hit of each gene in input blast file was used as recommended; options: cluster_type=collinear, alignment_method=gg2, max_gaps_in_alignment=10, tandem_gap=5, gap_size=10, cluster_gap=10, q_value=0.9, prob_cutoff=0.001, anchor_points=5, level_2_only=false), MCScanX (only top 5 hits of each gene in the input blast file was used as suggested; options: default) and SynChro (options: 0 6; 0 for all pairwise, and 6 for delta of RBH genes), for each program. To calculate synteny coverage, the total length of merged synteny blocks along scaffolds was divided by total assembly size.

### Data analysis

Visualization of synteny linkage was made by R (v 3.3.1) and circos [58] (v0.69-4). Gene ontology enrichment analysis was performed using topGO [59] (v1.0) package in R and only focused on Biological Process (options: nodeSize = 3, algorithm = “weight01”, statistic = “Fisher”). Gene ontology associations files for *C. elegans* and *C. briggsae* were downloaded from WormBase WS255 [24]. Gene orthology was assigned by OrthoFinder [57]. Then, assemblies were assembled using ALLMAPS [51] with a guided scaffolding approach. *de novo* Annotations of *C. elegans* was predicted using either the manually trained species parameter or from BUSCO [54] (v2.0).

## Acknowledgements

We thank John Wang for commenting the manuscript.

## Supporting information

**S1 Fig. Synteny coverage in different number of minimum anchor using DAGchainer. The** Y axis shows synteny coverage (%). The X axis is the number of minimum anchor to identify a synteny block from 2 to 8. The 4 colors are 4 combinations of synteny detection among species: *C. elegans* vs. *C. elegans* (CEvsCE, green), *C. elegans* vs. *C. briggsae* (CEvsCBG, orange), *S. ratti* vs. *S. ratti* (SRvsSR, blue) and *S.ratti* vs. *S. stercoralis* (SRvsSS, purple).

**S2 Fig. Synteny blocks in *C. elegans* versus *C. elegans.*** Chromosomes are separated into panels with Roman number labels. The Y axis stands for categories of distribution. Synteny blocks defined by four detection programs: DAGchainer (red), i-ADHoRe (yellow), MCScanX (green) and SynChro (light blue) are drawn as rectangles. Distribution of genes is the bottom smaller rectangles in burgundy. The X axis is the position of chromosome.

**S3 Fig. Synteny blocks in *C. elegans* versus 1Mb fragmented *C. elegans.*** Chromosomes are separated into panels with Roman number labels. The Y axis stands for categories of distribution. Synteny blocks defined by four detection programs: DAGchainer (red), i-ADHoRe (yellow), MCScanX (green) and SynChro (light blue) are drawn as rectangles. Distribution of genes is the bottom smaller rectangles in burgundy. The X axis is the position of chromosome.

**S4 Fig. Synteny blocks in *C. elegans* versus 100kb fragmented *C. elegans***.

Chromosomes are separated into panels with Roman number labels. The Y axis stands for categories of distribution. Synteny blocks defined by four detection programs: DAGchainer (red), i-ADHoRe (yellow), MCScanX (green) and SynChro (light blue) are drawn as rectangles. Distribution of genes is the bottom smaller rectangles in burgundy. The X axis is the position of chromosome.

**S5 Fig. A zoomed-in region of synteny identification with lower gap threshold in MCScanX between original *C. elegans* and 100kb fragmented assembly.** The Y axis stands for categories of distribution. Synteny blocks defined by four detection programs: DAGchainer (red), i-ADHoRe (yellow), MCScanX (green) and SynChro (light blue) are drawn as rectangles. Fragmented sites are labeled by vertical red dashed lines. Distribution of genes in burgundy rectangles is marked with dark blue lines as gene starts. The X axis is the position of chromosome. Scenario (a) is that synteny block was identified after gap threshold tuned lower.

**S6 Fig. Synteny blocks in *C. elegans* versus *C. briggsae.*** Chromosomes are separated into panels with Roman number labels. The Y axis stands for categories of distribution. Synteny blocks defined by four detection programs: DAGchainer (red), i-ADHoRe (yellow), MCScanX (green) and SynChro (light blue) are drawn as rectangles. The bottom four categories are orthologs between the two species assigned by Opscan (OP; burgundy) and OrthoFinder (OF; deep blue), and we further categorized orthologs into 1 to 1 orthology (1-1) or many to many orthology (N-N). The X axis is the position of chromosome.

**S1 Table. Statistics of annotation and synteny coverage using WormBase *C. elegans* versus PacBio *C. elegans***. Yellow highlights the statistics that relate to variation of annotation that may play a key role in synteny detections. Grey highlights the result of synteny detections by DAGchainer. Assembly source in column 2 is obtained from Pacific Bioscienceces Dataset. Annotation information in column 2 is predicted by Augustus using implanted caenorhabditis (elegans) species data set.

**S2 Table. Quantification of synteny coverage and error rate.**

**S3 Table. Gene ontology (GO) enrichment analysis of *C. briggsae* genes in synteny break between *C. elegans* and 1Mb fragmented *C. briggsae* assemblies.** GO terms that appeared in the top 10 ranks either in the original comparison or after when assemblies were fragmented, are displayed. The original rank, median rank and number of occurrences that reached top 10 in 100 replications are shown for each GO term. GO terms not belonging to original assembly but reached top 10 after fragmentation are shaded in green.

**S4 Table. Gene ontology (GO) enrichment analysis of *C. briggsae* genes in synteny break between *C. elegans* and 500kb fragmented *C. briggsae* assemblies.** GO terms that appeared in the top 10 ranks either in the original comparison or after when assemblies were fragmented, are displayed. The original rank, median rank and number of occurrences that reached top 10 in 100 replications are shown for each GO term. GO terms not belonging to original assembly but reached top 10 after fragmentation are shaded in green.

**S5 Table. Gene ontology (GO) enrichment analysis of *C. briggsae* genes in synteny break between *C. elegans* and 200kb fragmented *C. briggsae* assemblies.** GO terms that appeared in the top 10 ranks either in the original comparison or after when assemblies were fragmented, are displayed. The original rank, median rank and number of occurrences that reached top 10 in 100 replications are shown for each GO term. GO terms not belonging to original assembly but reached top 10 after fragmentation are shaded in green. GO:0043066 was in the original top 10 rank but failed to reach top 10 in all of 100 replications.

**S6 Table. Gene ontology (GO) enrichment analysis of *C. briggsae* genes in synteny break between *C. elegans* and 100kb fragmented *C. briggsae* assemblies.** GO terms that appeared in the top 10 ranks either in the original comparison or after when assemblies were fragmented, are displayed. The original rank, median rank and number of occurrences that reached top 10 in 100 replications are shown for each GO term. GO terms not belonging to original assembly but reached top 10 after fragmentation are shaded in green. GO:0043066 was in the original top 10 rank but failed to reach top 10 in all of 100 replications.

**S7 Table. Assembly statistics among *Caenorhabditis* species and *Strongyloides* species including ALLMAPS results.**

## References

1. Gordon D, Huddleston J, Chaisson MJ, Hill CM, Kronenberg ZN, Munson KM, et al. Long-read sequence assembly of the gorilla genome. Science. 2016;352: aae0344. doi:10.1126/science.aae0344

2. Lien S, Koop BF, Sandve SR, Miller JR, Matthew P, Leong JS, et al. The Atlantic salmon genome provides insights into rediploidization. Nature. 2016;533: 200–205. doi:10.1038/nature17164

3. Iorizzo M, Ellison S, Senalik D, Zeng P, Satapoomin P, Huang J, et al. A high-quality carrot genome assembly provides new insights into carotenoid accumulation and asterid genome evolution. Nat Genet. 2016;advance on: 657–666. doi:10.1038/ng.3565

4. Jarvis DE, Ho YS, Lightfoot DJ, Schmöckel SM, Li B, Borm TJA, et al. The genome of Chenopodium quinoa. Nature. 2017; 1–6. doi:10.1038/nature21370

5. Ma L, Chen Z, Huang DW, Kutty G, Ishihara M, Wang H, et al. Genome analysis of three Pneumocystis species reveals adaptation mechanisms to life exclusively in mammalian hosts. Nat Commun. Nature Publishing Group; 2016;7: 10740. doi:10.1038/ncomms 10740

6. de Man TJB, Stajich JE, Kubicek CP, Teiling C, Chenthamara K, Atanasova L, et al. Small genome of the fungus Escovopsis weberi, a specialized disease agent of ant agriculture. Proc Natl Acad Sci. 2016;113: 3567–3572. doi:10.1073/pnas.1518501113

7. Hunt VL, Tsai IJ, Coghlan A, Reid AJ, Holroyd N, Foth BJ, et al. The genomic basis of parasitism in the Strongyloides clade of nematodes. Nat Genet. Nature Publishing Group; 2016;48: 1–11. doi: 10.1038/ng.3495

8. Cotton JA, Bennuru S, Grote A, Harsha B, Tracey A, Beech R, et al. The genome of Onchocerca volvulus, agent of river blindness. Nat Microbiol. 2016;2: 16216. doi:10.1038/nmicrobiol.2016.216

9. Chen X, Tompa M. Comparative assessment of methods for aligning multiple genome sequences. Nat Biotechnol. Nature Publishing Group; 2010;28: 567–572. doi:10.1038/nbt.1637

10. Alkan C, Coe BP, Eichler EE. Genome structural variation discovery and genotyping. Nat Rev Genet. Nature Publishing Group; 2011;12: 363–76. doi:10.1038/nrg2958

11. Treangen TJ, Salzberg SL. Repetitive DNA and next-generation sequencing: computational challenges and solutions. Nat Rev Genet. 2012;46: 36–46. doi:10.1038/nrg3164

12. Uricaru R, Michotey C, Chiapello H, Rivals E. YOC, A new strategy for pairwise alignment of collinear genomes. BMC Bioinformatics. ???; 2015;16: 111. doi:10.1186/s12859-015-0530-3

13. Ehrlich J, Sankoff D, Nadeau JH. Synteny conservation and chromosome rearrangements during mammalian evolution. Genetics. 1997;147: 289–296. doi: 10.1159/000322358

14. Ghiurcuta CG, Moret BME. Evaluating synteny for improved comparative studies. Bioinformatics. 2014;30: 9–18. doi: 10.1093/bioinformatics/btu259

15. Renwick JH. The mapping of human chromosome. Annu Rev Genet. 1971;5: 81–120.

16. Nadeau JH. Maps of linkage and synteny homologies between mouse and man. Trends Genet. 1989; 1–5.

17. Vergara IA, Chen N. Large synteny blocks revealed between Caenorhabditis elegans and Caenorhabditis briggsae genomes using OrthoCluster. BMC Genomics. 2010;11: 516. doi:10.1186/1471-2164-11-516

18. Tang H, Lyons E, Pedersen B, Schnable JC, Paterson AH, Freeling M. Screening synteny blocks in pairwise genome comparisons through integer programming. 2011; 1–11.

19. Schmidt R. Synteny - Recent Advances and Future Prospects. Curr Opin Plant Biol. 2000;3: 97–102.

20. Vandepoele K, Saeys Y, Simillion C, Raes J, Van de Peer Y. The automatic detection of homologous regions (ADHoRe) and its application to microcolinearity between Arabidopsis and rice. Genome Res. 2002;12: 1792–1801. doi:10.1101/gr.400202

21. Coghlan A, Eichler EE, Oliver SG, Paterson AH, Stein L. Chromosome evolution in eukaryotes: A multi-kingdom perspective. Trends Genet. 2005;21: 673–682. doi:10.1016/j.tig.2005.09.009

22. Molinari NA, Petrov DA, Price HJ, Smith JD, Gold JR, Vassiliadis C, et al. Synteny and Collinearity in Plant Genomes. Science (80-). 2008; 486–489. Available: http://www.sciencemag.org/content/320/5875/486.full.pdf

23. Zhang G, Li B, Li C, Gilbert MTP, Jarvis ED, Wang J. Comparative genomic data of the Avian Phylogenomics Project. Gigascience. 2014;3: 26. doi: 10.1186/2047-217X-3-26

24. Howe KL, Bolt BJ, Cain S, Chan J, Chen WJ, Davis P, et al. WormBase 2016: Expanding to enable helminth genomic research. Nucleic Acids Res. 2016;44: D774–D780. doi:10.1093/nar/gkv1217

25. Consortium TC elegans S. Genome Sequence of the Nematode C. elegans: A Platform for Investigating Biology. Science (80-). 1998;282: 2012–2018. doi: 10.1126/science.282.5396.2012

26. Stein LD, Bao Z, Blasiar D, Blumenthal T, Brent MR, Chen N, et al. The genome sequence of Caenorhabditis briggsae: A platform for comparative genomics. PLoS Biol. 2003;1. doi:10.1371/journal.pbio.0000045

27. Wong S, Wolfe KH. Birth of a metabolic gene cluster in yeast by adaptive gene relocation. Nat Genet. 2005;37: 777–782. doi:10.1038/ng1584

28. Lemons D, McGinnis W. Genomic evolution of Hox gene clusters. Science (80-). 2006/09/30. 2006;313: 1918–1922. doi: 10.1126/science.1132040

29. Ruelens P, de Maagd RA, Proost S, Theißen G, Geuten K, Kaufmann K. FLOWERING LOCUS C in monocots and the tandem origin of angiosperm-specific MADS-box genes. Nat Commun. 2013;4: 2280. doi:10.1038/ncomms3280

30. Kemkemer C, Kohn M, Cooper DN, Froenicke L, Högel J, Hameister H, et al. Gene synteny comparisons between different vertebrates provide new insights into breakage and fusion events during mammalian karyotype evolution. BMC Evol Biol. 2009;9: 84. doi:10.1186/1471-2148-9-84

31. Murat F, Armero A, Pont C, Klopp C, Salse J. Reconstructing the genome of the most recent common ancestor of flowering plants. Nat Genet. Nature Publishing Group; 2017;49: 490–496. doi: 10.1038/ng.3813

32. Denton JF, Lugo-Martinez J, Tucker AE, Schrider DR, Warren WC, Hahn MW. Extensive Error in the Number of Genes Inferred from Draft Genome Assemblies. PLoS Comput Biol. 2014;10. doi:10.1371/journal.pcbi.1003998

33. Dupont P-Y, Cox MP. Genomic Data Quality Impacts Automated Detection of Lateral Gene Transfer in Fungi. G3 (Bethesda). 2017;7: g3.116.038448. doi:10.1534/g3.116.038448

34. Batzoglou S. The many faces of sequence alignment. Brief Bioinform. 2005;6: 6–22. doi: 10.1093/bib/6.1.6

35. Haas BJ, Delcher AL, Wortman JR, Salzberg SL. DAGchainer: A tool for mining segmental genome duplications and synteny. Bioinformatics. 2004;20: 3643–3646. doi:10.1093/bioinformatics/bth397

36. Wang Y, Tang H, Debarry JD, Tan X, Li J, Wang X, et al. MCScanX: A toolkit for detection and evolutionary analysis of gene synteny and collinearity. Nucleic Acids Res. 2012;40: 1–14. doi:10.1093/nar/gkr1293

37. Proost S, Fostier J, De Witte D, Dhoedt B, Demeester P, Van De Peer Y, et al. i-ADHoRe 3.0-fast and sensitive detection of genomic homology in extremely large data sets. Nucleic Acids Res. 2012;40: 1–11. doi: 10.1093/nar/gkr955

38. Drillon G, Carbone A, Fischer G. SynChro: A fast and easy tool to reconstruct and visualize synteny blocks along eukaryotic chromosomes. PLoS One. 2014;9: 1–8. doi:10.1371/journal.pone.0092621

39. Ross JA, Koboldt DC, Staisch JE, Chamberlin HM, Gupta BP, Miller RD, et al. Caenorhabditis briggsae recombinant inbred line genotypes reveal inter-strain incompatibility and the evolution of recombination. PLoS Genet. 2011;7. doi:10.1371/journal.pgen.1002174

40. Bhutkar A, Russo S, Smith TF, Gelbart WM. Techniques for multi-genome synteny analysis to overcome assembly limitations. Genome Inform. 2006;17: 152–161. Available: http://eutils.ncbi.nlm.nih.gov/entrez/eutils/elink.fcgi?dbfrom=pubmed&id=17503388&retmode=ref&cmd=prlinks%5Cnpapers3://publication/uuid/7717ABDA-5CCB-48C2-9AF7-C51B12BDEAF8

41. Goodwin S, McPherson JD, McCombie WR. Coming of age: ten years of next-generation sequencing technologies. Nat Rev Genet. Nature Publishing Group; 2016;17: 333–351. doi:10.1038/nrg.2016.49

42. Treangen TJ, Ondov BD, Koren S, Phillippy AM. The Harvest suite for rapid core-genome alignment and visualization of thousands of intraspecific microbial genomes. Genome Biol. 2014;15: 524. doi:10.1186/s13059-014-0524-x

43. Viney ME. The biology and genomics of Strongyloides. Med Microbiol Immunol. 2006;195: 49–54. doi:10.1007/s00430-006-0013-2

44. Ward JD. Rendering the intractable more tractable: Tools from caenorhabditis elegans ripe for import into parasitic nematodes. Genetics. 2015. pp. 1279–1294. doi:10.1534/genetics.115.182717

45. Armengol L, Marquès-Bonet T, Cheung J, Khaja R, González JR, Scherer SW, et al. Murine segmental duplications are hot spots for chromosome and gene evolution. Genomics. 2005;86: 692–700. doi:10.1016/j.ygeno.2005.08.008

46. Davidson RM, Gowda M, Moghe G, Lin H, Vaillancourt B, Shiu SH, et al. Comparative transcriptomics of three Poaceae species reveals patterns of gene expression evolution. Plant J. 2012;71: 492–502. doi:10.1111/j.1365-313 X.2012.05005.x

47. Lovell P V, Wirthlin M, Wilhelm L, Minx P, Lazar NH, Carbone L, et al. Conserved syntenic clusters of protein coding genes are missing in birds. Genome Biol. 2014; 1–27. doi:10.1186/s13059-014-0565-1

48. Baldauf J, Marcon C, Paschold A, Hochholdinger F. Nonsyntenic genes drive tissue-specific dynamics of differential, nonadditive and allelic expression patterns in maize hybrids. Plant Physiol. 2016;171: pp.00262.2016. doi:10.1104/pp.16.00262

49. Assefa S, Keane TM, Otto TD, Newbold C, Berriman M. ABACAS: Algorithm-based automatic contiguation of assembled sequences. Bioinformatics. 2009;25: 1968–1969. doi:10.1093/bioinformatics/btp347

50. Husemann P, Stoye J. r2cat: Synteny plots and comparative assembly. Bioinformatics. 2009;26: 570–571. doi: 10.1093/bioinformatics/btp690

51. Tang H, Zhang X, Miao C, Zhang J, Ming R, Schnable JC, et al. ALLMAPS: robust scaffold ordering based on multiple maps. Genome Biol. 2015;16: 3. doi:10.1186/s13059-014-0573-1

52. Howe KL, Bolt BJ, Cain S, Chan J, Chen WJ, Davis P, et al. WormBase 2016: Expanding to enable helminth genomic research. Nucleic Acids Res. 2016;44: D774–D780. doi:10.1093/nar/gkv1217

53. Stanke M, Steinkamp R, Waack S, Morgenstern B. AUGUSTUS: a web server for gene finding in eukaryotes. Nucleic Acids Res. 2004;32: W309–W312. doi: 10.1093/nar/gkh379

54. Simão FA, Waterhouse RM, Ioannidis P, Kriventseva E V. BUSCO : assessing genome assembly and annotation completeness with single-copy orthologs. Genome Anal. 2015;31: 9–10. doi:10.1093/bioinformatics/btv351

55. Hillier LW, Miller RD, Baird SE, Chinwalla A, Fulton LA, Koboldt DC, et al. Comparison of C. elegans and C. briggsae genome sequences reveals extensive conservation of chromosome organization and synteny. PLoS Biol. 2007;5: 1603–1616. doi:10.1371/journal.pbio.0050167

56. Chain PSG, Grafham D V, Fulton RS, Fitzgerald MG, Hostetler J, Muzny D, et al. Genome Project Standards in a New Era of Sequencing. Science. 2009;326: 4–5. doi: 10.1126/science. 1180614

57. Emms DM, Kelly S. OrthoFinder: solving fundamental biases in whole genome comparisons dramatically improves orthogroup inference accuracy. Genome Biol. Genome Biology; 2015;16: 157. doi:10.1186/s13059-015-0721-2

58. Krzywinski M, Schein J, Birol I, Connors J, Gascoyne R, Horsman D, et al. Circos: an information esthetic for comparative genomics. Genome Res. 2009;19: 1639–1645. doi:10.1101/gr.092759.109

59. Alexa A and Rahnenfuhrer J. topGO: Enrichment Analysis for Gene Ontology. In: R package version 2.26.0. [Internet]. 2016. Available: http://bioconductor.org/packages/release/bioc/html/topGO.html

